# A Novel Drosophila Model to Investigate Adipose Tissue Macrophage Infiltration (ATM) and Obesity highlights the Therapeutic Potential of Attenuating Eiger/TNFα Signaling to Ameliorate Insulin Resistance and ATM

**DOI:** 10.1101/2023.07.06.548016

**Authors:** Zhasmine Mirzoyan, Alice Valenza, Sheri Zola, Carola Bonfanti, Lorenzo Arnaboldi, Nicholas Ferrari, John Pollard, Valeria Lupi, Matteo Cassinelli, Matteo Frattaroli, Mehtap Sahin, Maria Enrica Pasini, Paola Bellosta

## Abstract

Obesity is a global health concern associated with various metabolic disorders including insulin resistance and adipose tissue inflammation characterized by adipose tissue macrophage (ATM) infiltration. In this study, we present a novel *Drosophila* model to investigate the mechanisms underlying ATM infiltration and its association with obesity-related pathologies. Furthermore, we demonstrate the therapeutic potential of attenuating Eiger/TNFα signaling to ameliorate insulin resistance and ATM. To study ATM infiltration and its consequences, we established a novel *Drosophila* model (OBL) that mimics key aspects of human adipose tissue and allows for investigating ATM infiltration and other related metabolic disorders in a controlled experimental system. We employed genetic manipulation to reduce ecdysone levels to prolong the larval stage. These animals are hyperphagic, and exhibit features resembling obesity in mammals, including increased lipid storage, adipocyte hypertrophy, and high levels of circulating glucose. Moreover, we observed a significant infiltration of immune cells (hemocytes) in the fat bodies accompanied by insulin resistance and systemic metabolic dysregulation. Furthermore, we found that attenuation of Eiger/TNFα signaling and using metformin and anti-oxidant bio-products like anthocyanins led to a reduction in ATM infiltration and improved insulin sensitivity.

Our data suggest that the key mechanisms that trigger immune cell infiltration into adipose tissue are evolutionarily conserved and may provide the opportunity to develop *Drosophila* models to better understand pathways critical for immune cell recruitment into adipose tissue, in relation to the development of insulin resistance in metabolic diseases such as obesity and type 2 diabetes, and non-alcoholic fatty liver disease (NAFLD). We believe that our OBL model can also be a valuable tool and provide a platform either to perform genetic screens or to test the efficacy and safety of novel therapeutic interventions for these diseases.

## INTRODUCTION

The dysregulation of fat metabolism affects organs and tissues, particularly the liver and the adipose tissue which regulate lipid homeostasis and triglycerides (Kahn et al., 2019; Morigny et al., 2021). The adipose tissue functions as an endocrine organ producing hormones and cytokines that, in pathological conditions such as obesity or metabolic syndrome, produces pro-inflammatory cytokines including tumor necrosis factor-alpha (TNFα) and interleukin-6 (IL-6), all of which can disrupt insulin signaling pathways and impair glucose uptake and metabolism, and also recruit immune cells, including macrophages, leading to chronic inflammation or adipose tissue macrophage infiltration (ATM) (Cai et al., 2022; Dahik et al., 2020; Gregor and Hotamisligil, 2011; Kawai et al., 2021). Furthermore, chronic inflammation is associated with a lifelong increase in oxidative stress and the production of high levels of reactive oxygen species (ROS) (Rani et al., 2016), often associated with the activation of JNK/SAPK/p46 signaling and apoptosis (Hotamisligil, 2003).

Obesity is associated with impaired fat metabolism, which results in elevated levels of circulating free fatty acids (FFAs) and post-translational modification of proteins by metabolites and lipids that can alter protein function. These have been shown to contribute to the development of insulin resistance, a major risk factor for various metabolic disorders, including non-alcoholic fatty liver disease (NAFLD) (Deprince et al., 2020; Yang et al., 2018), and increased susceptibility to cardiovascular disease (Nakamura and Sadoshima, 2020). Other important emergent regulators of fat metabolism are the Toll and IMD pathways, which are primarily associated with immune responses in both flies and humans (Leulier and Lemaitre, 2008; Yu et al., 2022), and have also been implicated in obesity and metabolic disorders. In humans Toll-like receptor TLR4 has been linked to obesity-related inflammation and is activated by fatty acids and lipopolysaccharides (LPS) derived from gut bacteria. This activation triggers a pro-inflammatory response, leading to the production of inflammatory cytokines and the activation of NF-κB, which can contribute to insulin resistance and metabolic dysfunction (Baker et al., 2011).

In recent years, a defined role for steroid hormones, such as ecdysone in insects (Kannangara et al., 2021) and estrogen and testosterone in mammals has been associated not only with their role in controlling the timing and progression of maturation but also with metabolic disorders, including the development of obesity. In particular, in humans, low levels of estrogen after menopause, or testosterone in hypogonadism, increases the risk of developing obesity or other metabolic complications (Fan et al., 2019; Sebo and Rodeheffer, 2021; Wittert and Grossmann, 2022) highlighting the remarkable similarities in the mechanisms underlying this regulation between evolutionarily distant species like flies and humans.

The *Drosophila* fat body (FB) is a metabolic tissue with similar physiological functions to the mammalian adipose tissue and liver that acts, in addition to storing nutrients and as an endocrine organ, to control key metabolic processes linked to the native immune response (Liu et al., 2009). In *Drosophila* the immune response is primarily mediated by the hemocytes or plasmacytes (macrophage-like), circulating in the hemolymph, present at all stages of the life cycle and compose the fly’s innate immune system (Buchon et al., 2014; Lemaitre and Hoffmann, 2007; Leulier and Lemaitre, 2008). Hemocytes are essential mediators in the cell-cell communication process: they have been shown to mediate a response between the FB to control the abnormal proliferation of tumor-like cells (Parisi et al., 2014) and to promote proliferation in response to ROS produced during cell death in the process of regeneration of the imaginal discs (Fogarty et al., 2016).

Studies in *Drosophila* have revealed the existence of conserved pathways that play a role in communication between metabolic organs. These signals are coordinated by secreted factors which allow organs to communicate with each other and coordinate various physiological processes (Droujinine and Perrimon, 2016). Among these, several adipokines are secreted by the FB and have important roles in metabolic regulation and communication between different tissues, including Eiger/TNFa that was shown to act on the insulin-producing cells (IPCs) to control the response to nutrients and to mediate the interplay between metabolism and immune signaling (Agrawal et al., 2016; Meschi and Delanoue, 2021). Additionally, Eiger/TNFα is known to be involved in inflammation and immune responses, and dysregulation of Eiger/TNF-α signaling has been associated with metabolic disorders such as insulin resistance and obesity (Igaki and Miura, 2014; Tzanavari et al., 2010; Yaribeygi et al., 2019).

By taking advantage of these conserved functional relationships between the immune cells (hemocytes) and the regulation of adipokines in the FB, we previously demonstrated that animals with reduced ecdysone have an obese-like phenotype and show high levels of hemocytes infiltrating the FB, mimicking what was described as chronic inflammation in human obesity. This was also accompanied by an increase in ROS and activation of the JNK/p46 signaling (Valenza et al., 2018).

Building upon this finding, we propose a model whereby blocking ecdysone could mimic conditions similar to the way reduction of estrogen in females, or testosterone in males, has the effect of regulating body fat distribution associated with greater risks of obesity-related metabolic disorders. Here we further characterized the metabolic changes in these obese larvae, which herein we refer to as OBL (obese larvae) and discovered that OBL larvae displayed hyperphagia leading to increased food intake. Because of this excessive feeding, the OBL larvae accumulated higher levels of triglycerides (TGAs), free fatty acids (FFAs), and glucose as they aged. These metabolic changes are consistent with what is often observed in obesity, where excess nutrients are stored as fat and circulating glucose levels are elevated. The cells of the FB in OBL are increased in size and store high levels of TAGs. Furthermore, the cells of cardiac tube show an accumulation of lipid-droplets and suffer from contraction-defects. The OBL animals do not respond to DILP2 regulation upon starvation, are hyperglycemic, and cells of their FBs have acquired insulin resistance. Furthermore, we found increased levels of Eiger/TNFa in the FB of the OBL larvae, and attenuation of Eiger signaling using animals heterozygous for *eiger^1^* or *eiger^3^* and their receptor *Grindelwald* (*grnd)* reduces the infiltration of hemocytes in the FB which also regains sensitivity to insulin treatment.

Finally, feeding the OBL animals with the anti-diabetic drug metformin and with natural antioxidant anthocyanins (ACN) significantly inhibits the infiltration of immune cells into the FB, confirming the possibility of using this new model to find novel drugs and to better understanding the programs that control the chronic inflammation and insulin resistance, in obesity and related metabolic disorders.

## RESULTS

### HFD and mutation in *brummer* lipases show low levels of hemocyte infiltration in the fat body

We previously demonstrated that reducing ecdysone in the prothoracic gland results in hyperphagic animals that accumulate fats and exhibit a low grade of hemocyte infiltration in the fat body (FB) (Valenza et al., 2018), resembling adipose tissue macrophage infiltration in humans or ATM (Kawai et al., 2021). To better visualize the hemocytes in our animals we used the hemolectin *Hml-DsRED* reporter line (Makhijani et al., 2011) and quantified the presence of hemocytes in the fat body of larvae either fed with a High-Fat Diet (HFD) (Diop et al., 2017) or in mutants for *brummer* lipase, the orthologue of human *PNPLA2/ATGL* lipase (Gronke et al., 2005). These data showed that feeding animals with an HFD significantly induced the migration of hemocytes in the FB of wt larvae, visible at 5 days after egg laying (AEL) (Figure 1A). In addition, this effect was further enhanced in animals mutant for Brummer lipase, suggesting that dysregulated lipid signaling or lipid accumulation favored the migration of the hemocytes in the FB. Using the *Hml-DsRED* hemocyte reporter line, we also re-analyzed and quantified the presence of the hemocytes in our animal model, which we called herein obesity-like model (OBL) genetically described in Supplementary Fig 1 (Valenza et al., 2018). In this OBL model, we observed a significant increase in the number of hemocytes infiltrating the FB as early as 5 days AEL, which is further potentiated at 12 days AEL (Figure 1B). This was also accompanied by a significant increase in apoptosis of FB cells (Figure 1C). Analysis in other tissues relevant to animal physiology showed that hemocytes preferentially infiltrate the fat body as few Hml-DsRED cells were detected in the brain, gut or other tissues (Supplementary Fig 2). Furthermore, an accurate image of the HML-Ds-RED distribution in the FB of the OBL animals at 12 days AEL revealed that hemocytes form the typical “crown” structures around the fat cells, a characteristic also observed in the adipose tissues of obese people with ATM (Figure 1 D-E). Similar structures were previously observed using anti-SPARC antibodies to visualize hemocytes in the FB of obese animals at 12 days AEL (Figure F–G) (Valenza et al., 2018). Furthermore, at higher magnification, *Drosophila* hemocytes show the typical jagged morphology like that exhibited by human macrophages (Fig H). In summary, our data indicate that in flies, a diet rich in fat or an alteration in lipid metabolism results in attracting the immune cell hemocytes (macrophage-like cells) into the adipose tissue, resulting in a low level of inflammation.

**Figure 1:**
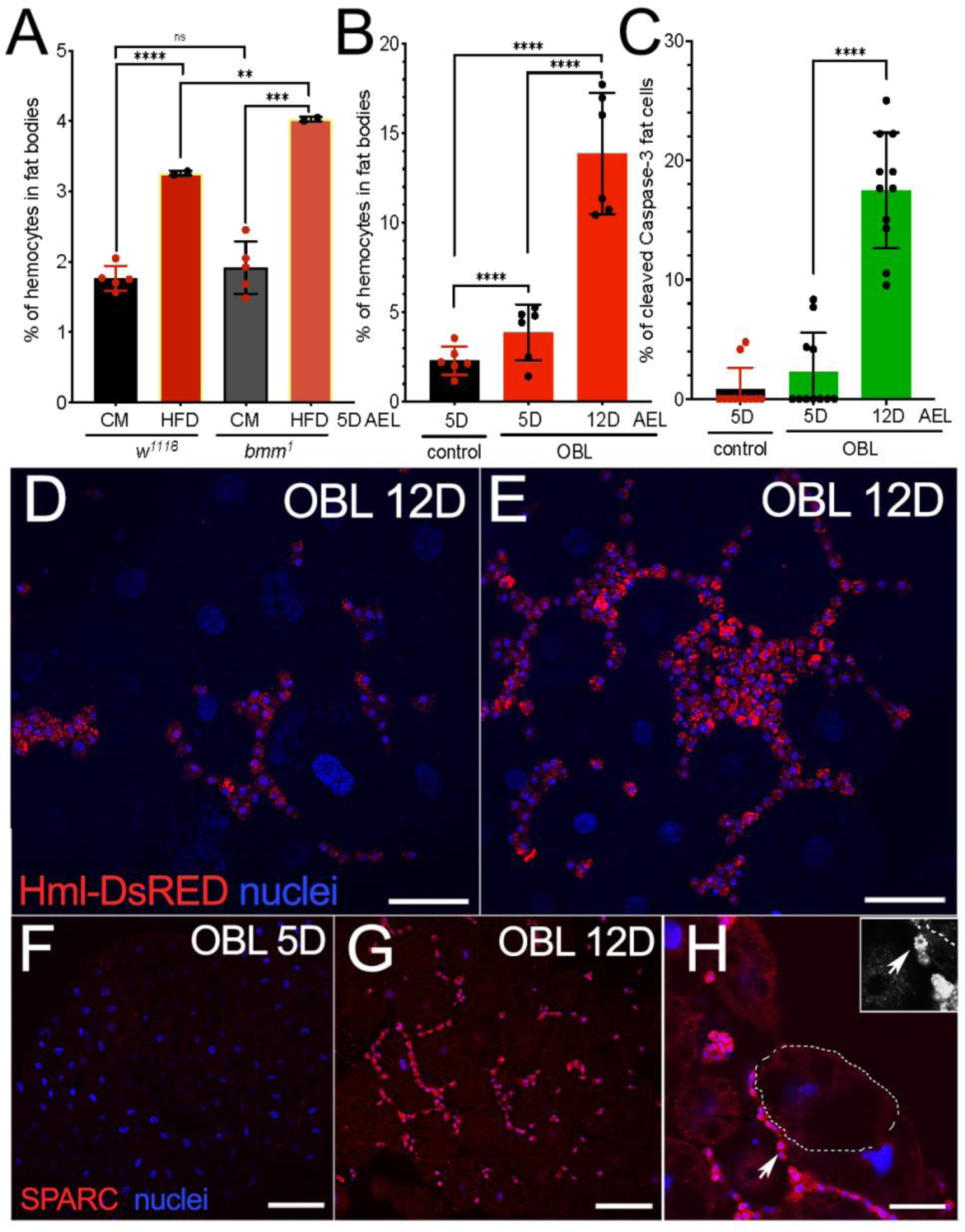
Hemocyte infiltration in the fat body of larvae fed with an HFD or mutants for *brummer* lipase, phenocopy low-grade chronic inflammation associated with obesity in humans. (A) Graphics representing the percentage of hemocytes infiltrating the FB of wild-type (*w^1118^)* animals fed with HFD or corn meal (CM) and in animals mutant for *brummer* lipase (*bmm^1^*) measured at 5 days AEL. (B) Quantification of the hemocytes in the FB measured at 5 days AEL in control *P0206>w^1118^*and in our model of obesity (OBL), at 5 and 12 days AEL. Quantification of hemocytes in the FB expressed as a percentage of the total number of cells visualized by nuclear staining using Hoechst. (C) Quantification of apoptosis in cells of the FB analyzed using anti-caspase3 antibodies in OBL animals at 12 days AEL (*p*<0.0001****). The scattered dots in A and B indicate the number of experiments (at least 10 animals were used for each time and genotype), and in C represents the number of animals used in the experiment. (D-E) Confocal images of FB from OBL animals at 12 days AEL showing the hemocytes marked by HML-Ds-RFP expression (Makhijani et al., 2011). (F-H) Confocal images of FB from *w^1118^* animals at 5 days (F) and of OBL at 12 days AEL (G and H) showing the hemocytes, stained using anti-SPARC antibodies (panel G is from (Valenza et al., 2018). In F is highlighted the crown-like structure formed by the hemocytes surrounding a fat cell. The islet shows a higher magnification of hemocytes (arrow) showing the typical macrophage-like morphology. The scale bar in panels D, E and H represents 50 μm, and in F-G represents 25 μm. Statistical analysis was performed using Student’s t-test and the error bars indicate the standard deviations. The asterisks represent the * = *p* < 0.05, ** = *p* < 0.01, *** = *p* < 0.001, and **** = *p* < 0.0001.

### Obese animals accumulate toxic TGAs and circulating FFAs, accompanied by lipid accumulation and contractility defects of the pericardial cells

Transcriptional analysis of key enzymes that regulate fat metabolism shows a decrease in *brummer-mRNA* expression in the obese-like larvae (OBL) at early larval development, while *Fatty Acid Synthase (FAS*) and *Perilipin-mRNAs* increase (Figure 2A-C). The OBL animals are hyperphagic and increase their size and weight over time (Fig 2D-E). Analysis of whole-body triglycerides (TAGs) shows that the OBL larvae accumulate TAG (Fig 2F) and increase their levels of Free Fatty Acids (FFAs) (Figure 2G). Lipidomic analysis (Fig 2H) confirms the significant increase in de-novo synthesis and accumulation of TAGs in OBL animals, as well as a significant increase in (SFA) saturated fatty acids (C:14 myristic C16 palmitic) over the Total lipids and of Monosaturated fatty acids (MUFA) (C16:1 and C18:1) (Figure 3) which in human pathologies such as obesity and T2D contribute to increased cellular fat and to the development of inflammation (Sarabhai et al., 2020; Silva Figueiredo et al., 2017; Wrzosek et al., 2022). Obese larvae also show a significant increase in TAG content, visualized by Nile red staining (Fig I), which parallels with the increase in the size of their fat cells (Figure 2J) and with the observed increase in TAGs and FFAs detected in the whole body.

**Figure 2:**
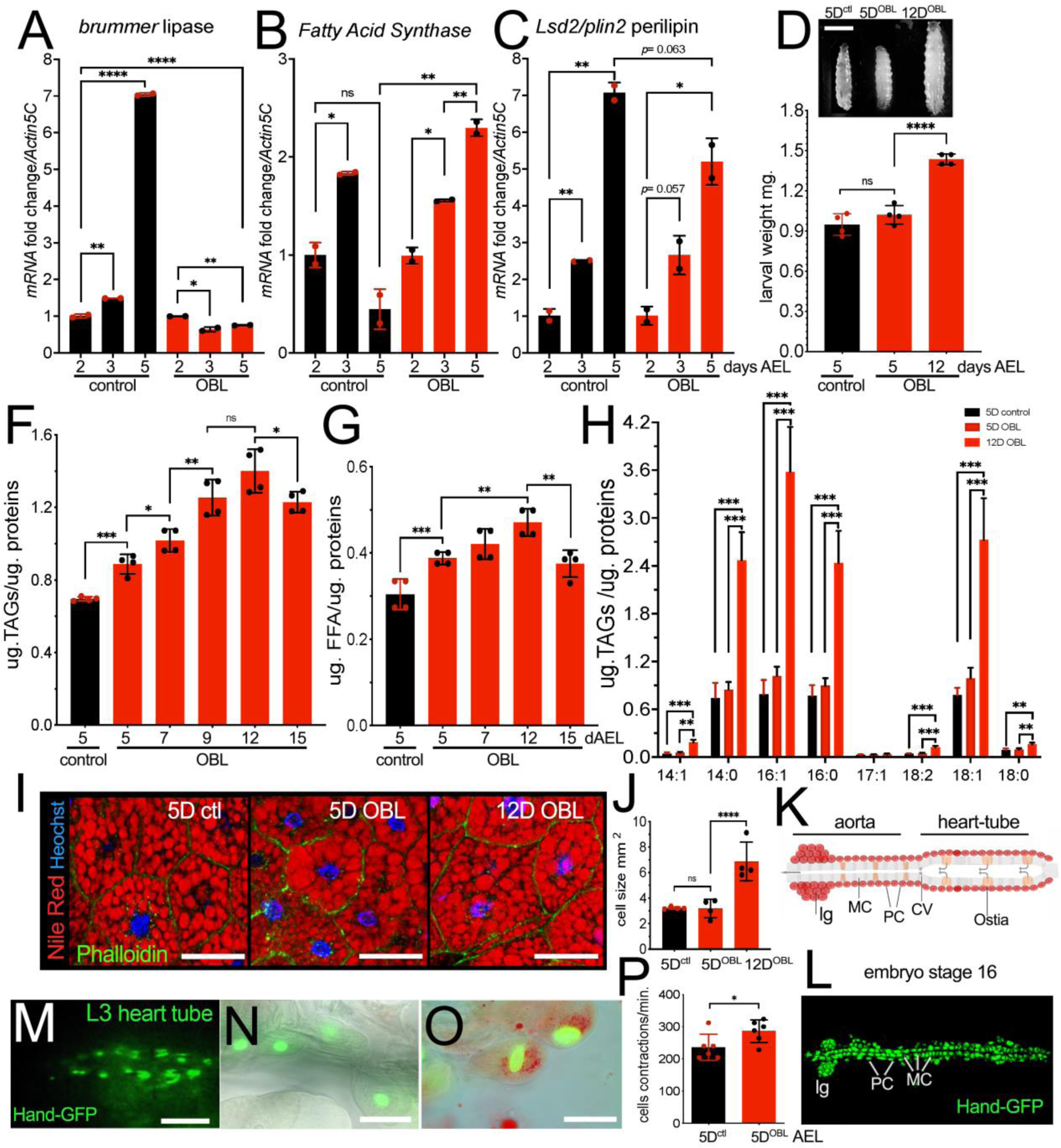
Obese animals increase TGAs and circulating FFAs, resulting in the accumulation of lipids in FB and pericardial cells. (A-C) qRT-PCT showing the level of *brummer, Fatty Acid Synthase*, and *Plin2-mRNAs* in whole larvae of the indicated genotype collected 2, 3, and 5 days AEL. (D) Photographs of control *w^1118^* or obese-like larvae (OBL) at 5 or 12 days AEL and (E) relative weight. (F) Content of triglycerides (TAGs) in whole larvae at the indicated time (days). (G) Free Fatty acids circulating in the hemolymph from larvae collected at the indicated time (days). (H) Lipidomic analysis of fats from whole larvae of the indicated genotype at 2, 5, and 12 days AEL. (I) Confocal images of cells from the FB of larvae of the indicated genotype, fat is stained with Nile red and nuclei with Hoechst. (J) Size-analysis of the cells from FB of animals at the indicated genotype, analysis was performed using ImageJ from confocal images. (K) Schematic draw of the cardiac tube in *Drosophila.* The heart tube contains openings called muscular ostia (Ostia) surrounded by muscle cells. The ostia function as valves to allow the entry of hemolymph into the heart tube. As the heart tube contracts, the hemolymph is propelled forward, filtered, and pushed out of the heart tube into the aorta, which is the main artery of the circulatory system in flies. The cardiovascular valve (CV) ensures the unidirectional flow of hemolymph from the heart tube to the aorta, preventing backflow. Myocardial Cells (MCs) and Pericardial Cells (PCs) are shown, lg is the lymph gland. (L) Photograph of a wild-type embryo (stage 16) showing the expression of Hand-GFP that mark the Myocardial Cells (MCs) and Pericardial Cells (PCs), expression is also detected in the lymph glands (lg). Anterior is to the left. (M) Photograph of the heart cardiac tube of a third instar larva where the pericardial cells are labeled with Hand-GFP. (N-O) Pericardial cells from third-instar larvae of the indicated genotype, marked by the expression of Hand-GFP and stained for lipid contents using Nile red. (P) Analysis of cardiac contraction in larvae at 5 days AEL of the indicated genotype. Scattered dots in the graphs indicate the number of experiments performed. The scale bar in panels: D represents 1 mm., in I represents 50 μm, in M 100 μm, and in N-O represents 20 μm. Statistical analysis was performed using Student’s t-test at exception for data in (H) where *p-*values were calculated from one-way analysis of variance (ANOVA) with Tukey multiple comparisons. The asterisks represent the * = *p* < 0.05, ** = *p* < 0.01, *** = *p* < 0.001, and **** = *p* < 0.0001, and the error bars indicate the standard deviations.

**Figure 3:**
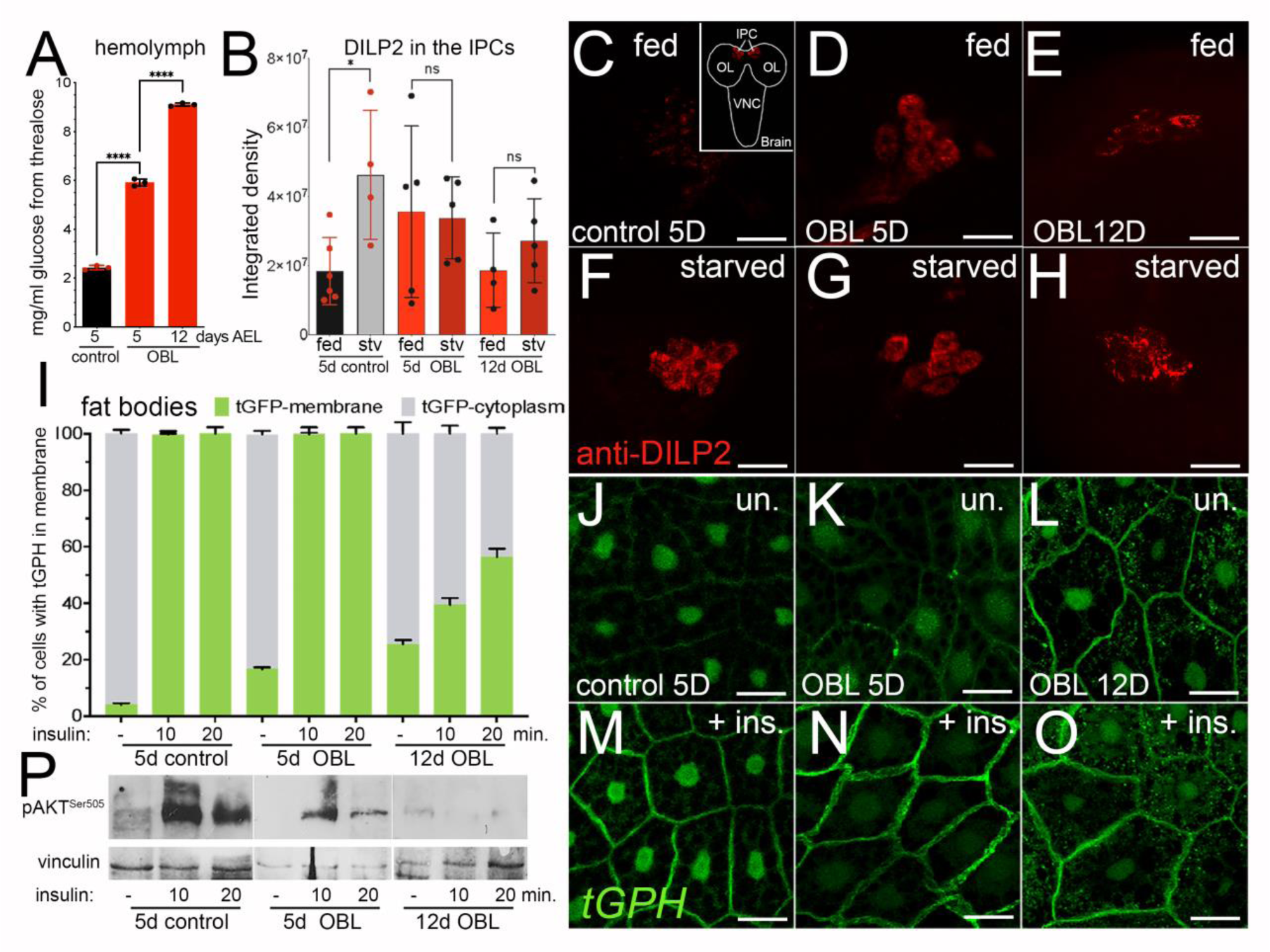
Obese larvae are hyperglycemic, have impaired systemic insulin signaling, and exhibit insulin resistance in the FB. (A) Hemolymph glucose analysis in animals of the indicated genotypes. At least 10 larvae were used for each genotype. (B) DILP2 was measured in the IPCs from third-instar larvae raised in corn meal food or starved for 24 hours in PBS using anti-DILP2 antibodies and the integrated density of fluorescence was quantified using confocal images as described previously (Parisi et al., 2013). *P* values were calculated from a Student t-test from n=10 larvae for each time point and genotype and from at least four independent experiments. Error bars represent the standard deviations. (C) Confocal images of brains from larvae of the indicated age and genotype raised in normal food (C-E) or in starvation (F-H) stained with anti-DILP2 and secondary antibody (red). (I) Quantification of tGPH (GFP) staining in the membrane of fat cells from dissected FB from larvae of the indicated genotypes treated ex-vivo with 1mM of insulin for the indicated time. (J-O) Confocal images of fat bodies from larvae expressing the tGPH reporter at the indicated age and genotype. (J-L) untreated (M-O) treated with insulin for 10 minutes. The scale bar in panels C-H and J-O represents 50 μm. (P) Western blot of lysates from fat bodies of animals of the indicated age and genotype upon treatment of insulin for the indicated time showing the level of phosphorylation of AKT-at Ser-505. Vinculin was used as a control.

Obesity is associated with cardiac dysfunction, therefore we examined whether there were any defects in the OBL larvae by analyzing the contraction of cardiomyocytes cells, visualized by the expression of Hand-GFP (Figure 2 M-P). Cardiac cells develop early in the embryo and are already contracting at stage 16 where the cardioblasts, which form the cardiac tube, differentiate into contractile cardiomyocytes or Myocardial Cells (MCs) while the Pericardial Cells (PCs) surrounds the MC cells to filter the hemolymph (Figure K and L) (Souidi and Jagla, 2021). Analysis of Hand-GFP cell contraction in the two genotypes shows an increase in the frequency of contraction in cells in the OBL larvae as early as 5 days AEL (Figure 2P), furthermore, these cells show an accumulation of lipid droplets (Fig 2N-O), suggesting that lipid dysregulation in the OBL larvae leads to heart defects similar to those induced by HFD (high fat) and HSD (high sugar) diets known to induce cardiomyopathies (Birse et al., 2010; Na et al., 2013).

### Obese animals have impaired insulin homeostasis and suffer from insulin resistance

Hyperglycemia is often associated with metabolic dysfunctions, such as obesity and type 2 diabetes (T2D), and is often accompanied by insulin resistance, particularly in metabolic tissues like adipocytes (Tabak et al., 2012). Previous work in flies showed that animals in HSD and HFD suffer from hyperglycemia and insulin resistance (Birse et al., 2010; Morris et al., 2012; Musselman et al., 2011; Skorupa et al., 2008). Our analysis shows that the OBL larvae are hyperglycemic and show an increase in circulating glucose levels (in the form of trehalose) already significant at 5 days AEL (Figure 3A), favoring the increase of glucose and storage of glycogen in the whole body (Supplementary Fig 4). Elevated glucose in the hemolymph has been associated with defects in the activity of *Drosophila* insulin-like peptides (DILPs), in particular of DILP2 and DILP5 (Semaniuk et al., 2018; Zhang et al., 2009). DILP2 secretion is controlled by the FB, which senses the levels of nutrients in the hemolymph to remotely control secreted factors that control DILP2 secretion from the brain (Geminard et al., 2009; Meschi and Delanoue, 2021). Analysis of DILP2 secretion shows that at 5 days AEL, while the IPCs of wt animals exhibited undetectable levels of DILP2 during normal feeding conditions, and accumulated DILP2 during starvation, OBL larvae show higher levels of DILP2 in IPCs in feeding and starved conditions both at 5 and 12 days AEL (Fig 3B and C-H), indicating that the FB is unable to control DILP2 homeostasis and it does not respond properly to the nutrient levels circulating in the hemolymph.

Excess circulating lipids lead to insulin resistance in adipose cells (Sarabhai et al., 2020), therefore we next analyzed if the cells of the FB were insensitive to insulin stimulation. In ex-vivo experiments, we isolated the fat bodies from control larvae and OBL larvae at 5 or 12 days AEL and incubated them with medium containing 1 mM insulin for the indicated times. Activation of insulin signaling was analyzed using the reporter line tGFP carrying the PH (pleckstrin homology) domain fused with green-fluorescent protein that translocates to the plasma membrane upon PI3K activation (Britton et al., 2002). These data showed that treating FBs from both control and OBL animals at 5-day AEL with insulin for 10 min completely induced the translocation of the tGFP reporter into the membrane (Figure 3I and J-O), whereas in FBs from 12-day OBL, insulin activity was reduced, with only 40% of the tGPH reporter translocated in the membrane, and the signal did not increase after 20 min of treatment (Figure I and J-O). Furthermore, we observed a low background level of tGPH signaling in the membrane of OBL animals at 12-day AEL even before the addition of insulin (Figure 3 L and O). We then analyzed how FB cells respond to insulin stimulation by measuring Akt phosphorylation on Ser-505 in fat lysates from control and OBL animals collected at 5 and 12 days AEL. These data show that while in control animals insulin treatment substantially activates AKT phosphorylation with a signal that lasted at least 20 minutes, in lysates from OBL larvae, the treatment was significantly reduced, and almost undetectable in OBL animals at 12 days AEL (Figure P). Together these analyses indicate that obese animals at 12 days AEL have impaired systemic insulin signals that result in fat bodies exhibiting insulin resistance.

### Eiger/TNFα signaling contributes to ROS production, chronic inflammation, and insulin resistance in the FB of obese animals

Inflammatory cytokines, like members of the TNFα family, are known to contribute to the low chronic inflammation described in ATM and other human metabolic disorders (Ferrante, 2007). Using the reporter line Eiger-GFP (Destefanis et al., 2022), we analyzed the level of Eiger, the sole *Drosophila* homolog of human TNFα (Igaki and Miura, 2014) in our OBL model. These data show that Eiger-GFP increased over time in the FB of the OBL animals (Figure 4A-C and graph in D), particularly visible in small cytoplasmic vesicles in the cells of the OBL larvae at 12 days AEL (Figure 4C), and this was accompanied by an increase in *eiger-mRNA* (Supplementary Fig 5). We then quantified the expression of Eiger-GFP in the hemocytes infiltrating the FBs and found that its expression was higher in the hemocytes of 12-day AEL larvae (Figure 4E-G and graph in H). We also noted that hemocytes were more abundant near FB cells with a higher density of Eiger-GFP vesicles, suggesting that non-autonomous Eiger secretion by the FB might function as an attractant for immune cells, which are responsible for engulfing and degrading dysfunctional fat cells.

**Figure 4.**
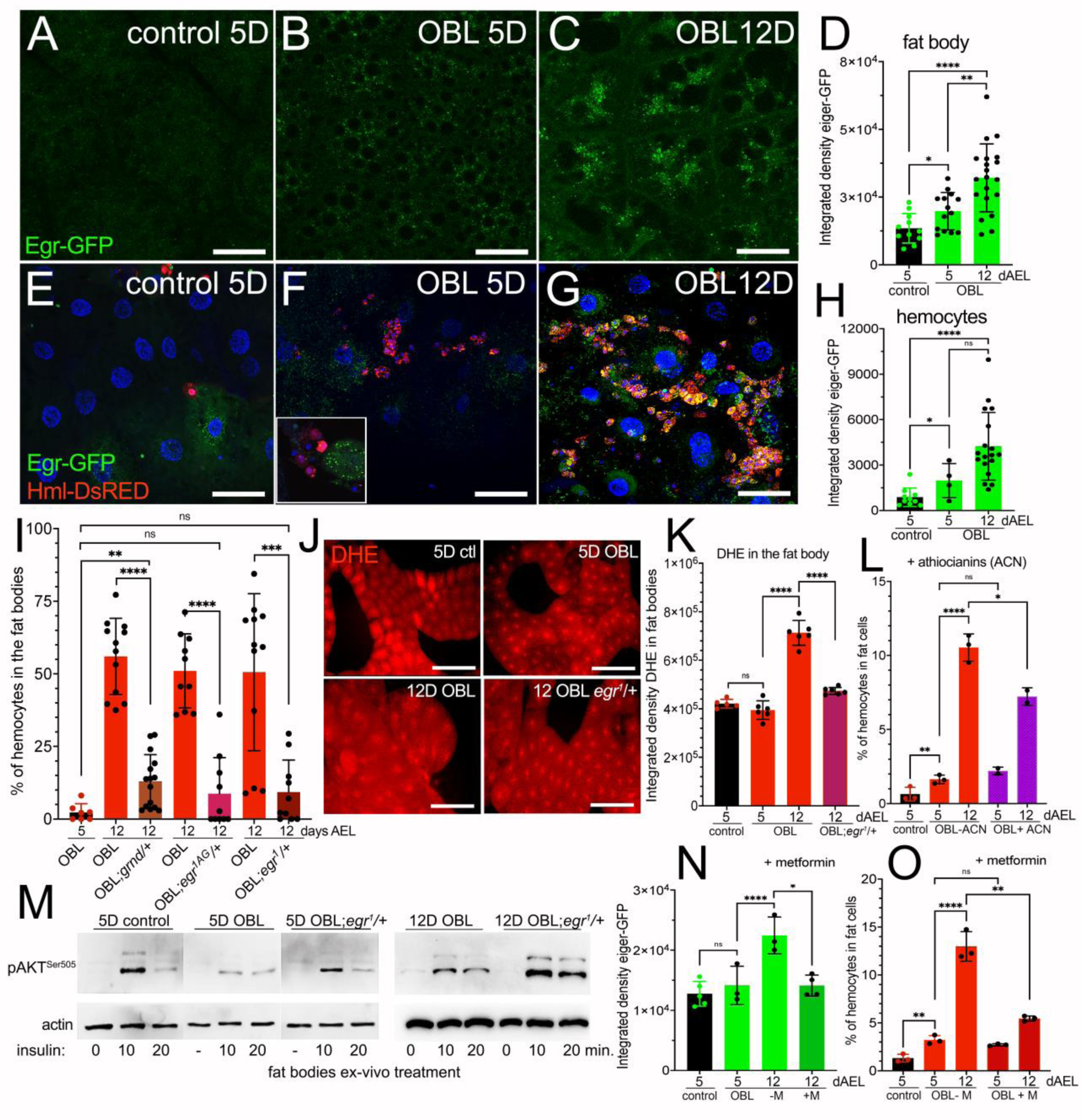
TNFa/eiger signaling in the FB and hemocytes of OBL animals contributes to the infiltration of hemocytes, ROS production, and insulin resistance. (A-C) Confocal images showing eiger-GFP expression in the FB from animals of the indicated genotype. (D) Quantification of the expression of eiger-GFP measured as integrated density in the FB of animals at the indicated time and genotype. (E-G) confocal images of FB showing the presence of hemocytes, labeled with Hml-Ds-RFP and of eiger-GFP. The inset in panel F highlights hemocytes near a FB cell expressing a high level of eiger-GFP. In panel G aggregates of hemocytes (red) are visible in OBL animals of 12 days AEL surrounding cells of the FB co-expressing eiger-GFP. (H) Quantification of eiger-GFP in hemocytes measured as integrated density in the FB from animals at the indicated time. The scale bar in panels A-C and D-F, represents 50 μm (I) Analysis and quantification of hemocytes infiltration in the FB from WT animals at 5 days AEL or in OBL animals heterozygous for *egr*^1^, *egr^1AG^* or *grnd* at 12 days AEL (K) DHE staining in FB and its quantification (J) measured as integrated density from animals of the indicated genotype. (K) Quantification of hemocytes infiltrating the FB in animals treated with anthocyanins (ACN). (K) Western blot using lysates from dissected fat bodies from animals of the indicated genotypes showing the level of AKT^S505^ phosphorylation upon treatment with insulin for the indicated time. Actin was used as a control for loading, western blot analysis was repeated twice using 20 fat bodies for each time point. (L-M) Quantification of eiger-GFP expression (L) or hemocytes (M) in FB after administration with the food of 1mM of metformin. *P* values were calculated from a Student t-test from n=10 larvae for each time point and genotype.

We thus tested the effect of reducing eiger signaling on the migration of hemocytes in the FB using mutants of e*iger* (*egr^1^, egr^3^*) (Igaki et al., 2002) or (*egr^1AG^, egr^3AG^*) (Kodra et al., 2020) and of the receptor *Grindelwald* (*grnd*) encoding for a transmembrane protein with homology to the tumor necrosis factor receptor (TNFR). (Andersen et al., 2015). In these experiments, we observed a dominant effect of these mutations that were all able to significantly reduce the migration of the hemocytes into the FB (Fig 4I). Moreover, our analyses showed similar results using *grnd* mutant and either *eiger^1^* and *egr^1A^*^G^ or *egr^3^* mutants, excluding the possibility of a functional effect of the NimroidC1 (NimC1) receptor, which was found to be deleted in the *egr^1^ and egr^3^* mutant lines (Kodra et al., 2020). We thus continued our analysis using *egr^1^* mutants and showed that a heterozygous mutant background for *egr^1^* significantly inhibits ROS production, measured with DHE staining in animals at 12 days AEL (Figure 4J). These data were corroborated by experiments in which the administration of the antioxidant anthocyanin (ACN) in food significantly reduced the infiltration of hemocytes into the FB of OBL animals at 12 days AEL (Fig 4K), and inhibited JNK/p46 phosphorylation, as we previously demonstrated in similar experiments using the OBL obesity model (Valenza et al., 2018). Furthermore, we show that in the FB of 5 and 12-day OBL animals heterozygous for *egr^1^*, AKT phosphorylation on Ser-505 was significantly recovered, suggesting that *eige*r heterozygous condition, improves the activity of insulin in the fats of our OBL animals (Figure 4K). Finally, we showed that the anti-diabetic drug metformin inhibits both, eiger-GFP upregulation (Figure 4L) and hemocyte infiltration in the FB of OBL animals at 12 days AEL (Figure 4 M). Overall, these data indicate that the mechanisms driving insulin resistance are conserved in our model of obesity, suggesting that it could be used either to screen for antidiabetic or anti-inflammatory drugs, like metformin, or to perform genetic screens to identify components that contribute to metabolic diseases such as obesity, T2D, and NAFLD.

### IMD and TOLL pathways contribute to the mechanism of hemocytes-infiltration in the FB of the obese larvae

Another mechanism that induces hemocyte activation in defense against infections and bacteria, is driven by the Toll and IMD pathways, in which Spaetzle (Spz), the ligand of the Toll receptor, initiates the humoral response in the FB to drive the secretion of antimicrobial peptides, including Diptericin (Dpt) and Drosomycin (Drs)(Lemaitre and Hoffmann, 2007). We found that the FB of OBL animals at 12 days AEL, activate the reporter lines Dpt-LacZ and Drs-GFP (Figure 5A). Furthermore, in a *spaetzle* mutant background, using the *spzr^m7^* allele (Morisato and Anderson, 1994) we saw that the number of hemocytes in the FB is significantly reduced (Figure 5B). Similar results were obtained using the *Rel^E20^* allele (Figure 5C), which carries a mutation for the gene *relish* (Hedengren et al., 1999) the close homolog of human NF-kb that acts downstream of both the IMD and Toll pathways (Lemaitre and Hoffmann, 2007). These data suggest that both cellular and humoral immunity cooperate to activate defense mechanisms, both cell-mediated and secreted, that are induced when FB cells and their lipid signaling are dysregulated (obesity) to attract hemocytes (macrophage-like cells) and eliminate them. *P* values were calculated from a Student t-test from n=10 larvae for each time point and genotype.

**Figure 5.**
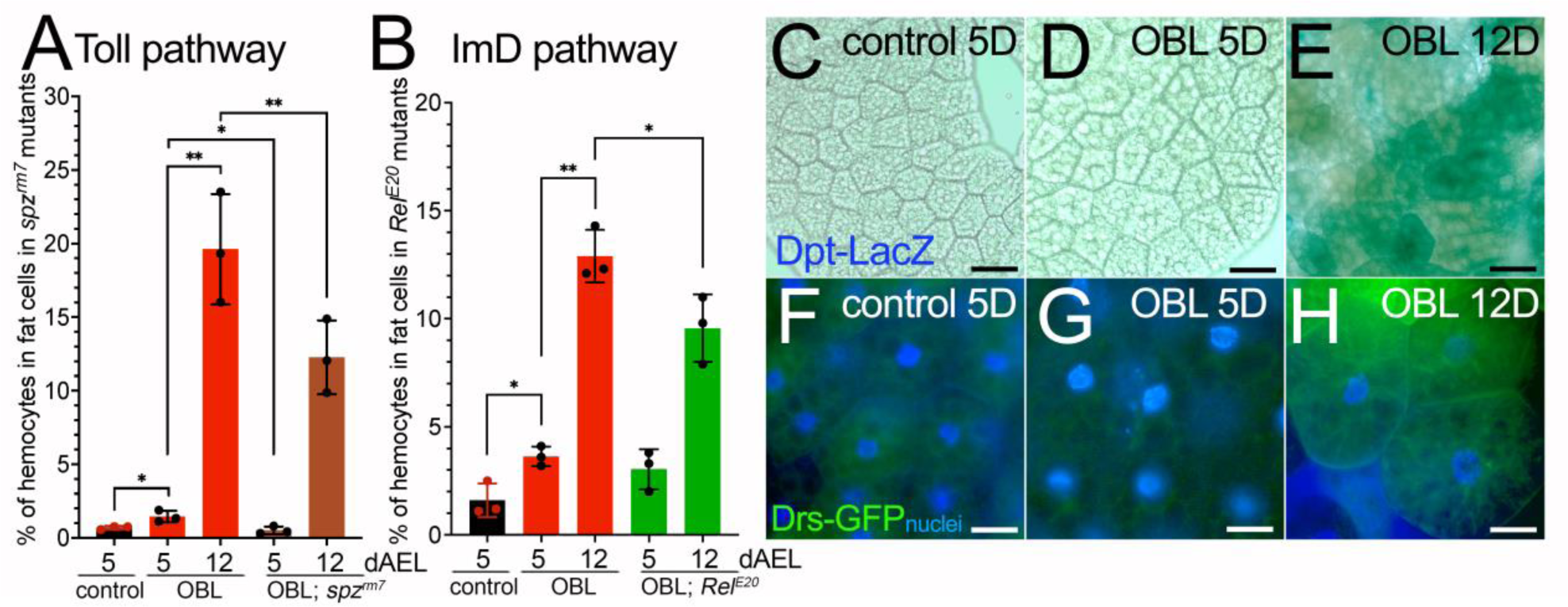
Spaetzle and Relish are involved in the mechanisms controlling hemocyte infiltration in the FB in the OBL animals. (A) Quantification of the hemocytes infiltrating the FB in WT animals at 5 days AEL or in OBL animals at 5 and 12 days AEL (red) and OBL; mutant for *Spaztle* carrying the *spzr^m7^* allele (brown). (B) Quantification of the hemocytes infiltrating the FB from WT of OBL and OBL carrying a mutation in *relish* using the *Rel^E20^* allele. (C-E) Confocal images of FB showing the reporter for Diptericin *Dpt-lacZ* staining by immunofluorescence using anti-beta-Gal antibodies in FB from WT or OBL animals at the indicated time of development. (F-H) Confocal images of FB showing the expression of Drosomycin using the *Drs-GFP* reporter line from WT or OBL animals at the indicated time of development. Nuclei are stained with Hoechst. The scale bar in panels C-E represents 20 μm, F-H represents 50 μm, in panels of J represents 20 μm. The scattered dots in graph A and B indicate the number of independent experiments (at least 10 animals were used for each time and genotype). Statistical analysis was performed using Student’s t-test and the error bars indicate the standard deviations. The asterisks represent the * = *p* < 0.05, ** = *p* < 0.01.

## DISCUSSION

### Hormonal regulation

Steroid hormones, like ecdysone in flies and estrogen/testosterone in humans, are involved in controlling the timing and progression of maturation. These hormones are synthesized in specific glands (prothoracic gland in flies and gonads in mammals) and released into the bloodstream to act on target tissues. They bind to specific receptors and initiate signaling cascades that result in the onset of the maturation processes. Similarly, insulin signaling pathways play a crucial role in regulating growth in both flies and mammals. In flies, insulin-like peptides (Dilps) secreted by insulin-producing cells (IPCs) in the brain bind to insulin receptors and activate insulin and insulin-like growth factor signaling (IIS) to promote growth and development (Buhler et al., 2018). The conservation of these mechanisms suggests that there are fundamental similarities in the ways organisms coordinate growth and maturation, regardless of their evolutionary distance. Steroid hormones and insulin signaling pathways have been evolutionarily conserved due to their important roles in regulating crucial developmental processes. In *Drosophila*, ecdysone triggers the transition from larval stages to pupal and adult stages (Kannangara et al., 2021). Similarly, in humans, steroid hormones are essential for the onset of puberty and the progression of sexual maturation. In addition to their roles in growth and maturation, steroid hormones such as estrogen and testosterone in mammals have also been implicated in metabolic regulation, including the development of obesity. By disrupting the steroid hormone ecdysone, we have generated a new model of obesity in flies that allowed us to further characterize obesity and the pathways involved in the development of insulin resistance, changes in fat metabolism, chronic inflammation, immune system dysfunction, and cardiac defects.

### Lipid metabolism

In obesity, adipose tissue undergoes hypertrophy and becomes dysfunctional. Our OBL animals are hyperphagic and like obese animals consume more food that accumulates as triglycerides within the adipose tissue and in the whole body (Figure 2). This adipose tissue dysfunction leads to an imbalance between lipolysis and lipogenesis, indeed in the OBL animals we observed an increase in FAS expression and in elevated levels of FFAs in the whole body.

We found in the OBL animals that Fatty Acid Synthase (FAS) is upregulated, a condition that is often found in human obesity and induces “de novo” synthesis of fatty acids, occurring by converting acetyl-CoA into long-chain fatty acids. Our lipidomic analysis confirms this situation since it shows a significant increase in de-novo synthesis and accumulation of TAGs in OBL animals, as well as a significant increase in (SFA) saturated fatty acids (C:14 myristic C16 palmitic) over the total lipids and of monosaturated fatty acids (MUFA) (C16:1 and C18:1) (Figure 2H), which in human pathologies such as obesity and T2D contribute to increased cellular fat and to the development of inflammation (Sarabhai et al., 2020; Silva Figueiredo et al., 2017; Wrzosek et al., 2022).

“de novo” synthesis of fatty acids can be induced by multiple factors such as insulin resistance, or by high levels of circulating insulin (Kojta et al., 2020). Indeed, the lack of response to insulin seen in the cells of the FB from the OBL animals (insulin resistance), and the upregulation of FAS may be related to the dysregulation of insulin singling, as shown in Figure 3I where higher levels of the tGPH reporter are visible in the membrane of the OBL animals and to the lack of control of DILPs regulation from the cells of the FB as seen by the abnormal accumulation of DILP2 in the IPC cells in fed and starved conditions (Figure 3B). Because of the systemic lipid imbalance, the excess of FFAs released by adipose tissue can be taken up by various organs (dyslipidemia), specifically we observed an accumulation of lipid droplet formations within the pericardial cells of the tubular heart (Figure 2N-O). This accumulation of lipids in pericardial cells can disrupt cellular processes and organelle function within pericardial cells, leading to impaired contractility of the heart muscle. This can manifest as contractility defects, reduced cardiac output, and potentially contribute to the development of cardiovascular complications in obese animals. Furthermore, this process of dyslipidemia, which we found induced when the FB is not functioning properly (Destefanis et al., 2022), may be responsible for the accumulation of lipids in other organs, such as the gut and brain (not shown), a condition that in humans is associated metabolic disorders such as obesity and type 2 diabetes (Matsuda and Shimomura, 2013).

In humans, secreted pro-inflammatory adipokines such as leptin (Friedman, 2019), which in *Drosophila* is functionally equivalent to Upd2 (Rajan and Perrimon, 2012), and TNFα are responsible for FAS expression and activation; similarly we found an upregulation of Eiger in the FB of OBL (Figure 4 and Supplementary 5), highlighting another conserved mechanism that can be further explored using our model to better understand the control of immune cell infiltration in the FB/adipocytes, metabolic disorders, and related pathologies. The study highlights the contribution of lipases, particularly ATGL (adipose triglyceride lipase), the human homologue of Brummer lipase, in the model of ATM (adipose tissue macrophage) and emphasizes the role of ATGL in this pathology. Previous research in flies has demonstrated that mutations in the *brummer* gene or alterations in its expression can affect lipid metabolism and fat storage in Drosophila (Gronke et al., 2005). We found that mutation of Brummer lipase favors hemocyte migration in the FB (Figure 1A). We speculate that dysregulated lipid metabolism resulting from dysfunctional lipases could lead to the production of toxic lipids, such as C16:0 palmitic acid and C16:1 palmitoleic acid. This dysregulation may be responsible for the increased ratio of saturated fatty acids (SFAs) to total lipids, as indicated in Supplementary Figure 3 and Figure 2H. The elevated ratio of SFAs to total lipids could then enhance signaling pathways that attract macrophages/hemocytes to migrate into the fat body.

A note should be mentioned about Brummer lipase since in *Drosophila* its activity and expression are under the control of the hormone ecdysone, which we reduced to generate our hyperphagic state, (Supplementary Figure 1). It is also important to note that the regulation of ATGL in obesity is complex and can vary depending on individual factors and the specific stage of obesity, and its activity is also influenced by growth hormone, and glucocorticoids (Patel et al., 2022). While in some cases an upregulation of ATGL can occur, there are instances where ATGL activity is impaired due to factors like chronic inflammation or dysregulated hormonal signaling, which is more similar to our OBL model. Nevertheless, the understanding of the role of lipases in obesity is still evolving. It is unclear whether directed mutations or reductions in Brummer/ATGL-like lipases contribute significantly to obesity, however, variations in genes related to lipid metabolism, including lipases, largely contribute to differences in obesity sensitivity, and reducing Brummer lipase activity in *Drosophila* may provide insights into the regulation of lipid metabolism in human obesity.

### Glucose regulation

Impaired insulin homeostasis is common in obesity and metabolic disorders which may lead to the development of insulin resistance, a state that contributes to the development of T2D. We found that OBL larvae exhibited insulin resistance in the FB (Figure 3I). It is important to note that while the term “insulin resistance” may not be applicable to *Drosophila* in the same sense as in mammals, here we have identified a condition in which the *Drosophila* FB exhibits similarities to insulin resistance seen in mammals. The cells of the FB in OBL animals are hypertrophic and release increased amounts of the pro-inflammatory cytokine Eiger (Figure 4). In addition, these animals show high levels of FFAs, conditions that in humans interfere with the function of insulin signaling in target tissues such as liver, muscle, and adipose tissue, contributing to insulin resistance.

Prolonged exposure to high insulin levels in obesity can lead to downregulation of insulin receptor on target cells, reducing their sensitivity to insulin and impairing downstream phosphorylation events. Here we show that cells of the FB in the OBL animals at 12 days AEL have a significant reduction of AKT phosphorylation. These mechanisms involve dysregulation of insulin-like signaling and impaired metabolic homeostasis. Previous studies in flies have shown that genetic manipulations of insulin signaling components, such as the insulin receptor (InR), can disrupt metabolic homeostasis and lead to abnormal glucose and lipid metabolism in the fat body (Baker and Thummel, 2007). Here we have mimicked a nutrient overload situation where hyperphagia and excessive food intake induce metabolic dysregulation and impair insulin-like signaling, leading to altered energy metabolism and nutrient storage. These conditions result in systemic metabolic imbalances, including “hyperglycemia” and aberrant glucose metabolism with the accumulation of glycogen and glucose (Supplementary Figure 4).

We want to also highlight how our model could be applicable for studying the metabolic alterations inducing NAFLD. Indeed, recent data show that NAFLD is not only a consequence of insulin resistance, but it is also an important cause of insulin resistance in the liver (Stefan et al., 2023). An important aspect of the pathologies mentioned above is their link to aging and cellular senescence (CS) (Spinelli et al., 2023) that was recently associated with insulin resistance, a common condition found in obesity, T2D, and NAFLD (Stefan et al., 2023).

While the specific molecular mechanisms underlying insulin-like signaling impairment and metabolic dysregulation are still being elucidated, studies using our OBL model could provide valuable insights into the regulation of metabolism and the potential factors contributing to metabolic dysfunction, including features that parallel aspects of insulin resistance seen in mammals.

### The Immune System and Inflammation-potential roles for Eiger/TNFα, IMD, and TOLL pathways

Studies in flies have shown that Eiger is released by the hemocytes upon stress conditions to produce ROS resulting in the activation of the JNK signaling pathway and apoptosis, suggesting the existence of non-autonomous signals between the hemocytes and the neighboring cells (Fogarty et al., 2016; Igaki and Miura, 2014; Parisi et al., 2014). In human obesity, the levels of TNFα increase in the adipose tissue activating pro-inflammatory cytokines (Ferrante, 2007). In a similar way, we found increased Eiger expression in the cells of the FB from OBL animals (Figure 4D); we hypothesize that this is fueling low-grade chronic inflammation by attracting hemocytes that migrate into the FB (Figure 4G). This data is accompanied by the transcriptional upregulation of *Glutathione S-transferase-D1* (*Gst-D1)* (Supplementary Figure 5), an enzyme that is usually upregulated to trigger cellular defense mechanisms and counteract the oxidative stress response in response to the activation of the JNK/p46 pathway (Khoshnood et al., 2016), a signaling that we previously found upregulated the FB of OBL animals at 12 days AEL (Valenza 2018). Reducing Eiger/TNF-α signaling, reduces the infiltration of the hemocytes in the FB (Figure 4I) and decreases ROS production in OBL fats (Figure 4J). Both chronic inflammation and elevated ROS levels contribute to insulin resistance, and we observed that OBL animals heterozygous for *egr^1^/*+ improved their response to insulin and showed a significant increase in AKT phosphorylation after ex-vivo treatment of their FBs with insulin (Figure 4M). Thus, we believe that the overall increased production of TNFα in adipose tissue is also induced in the OBL animals, causing the chronic inflammation and metabolic dysfunction observed in our animals and in obesity.

We know that in humans pro-inflammatory cytokines and factors contribute to the inflammatory state in obesity, indicating the complexity of the obesity-associated inflammatory response; however, we believe that the availability of a simple and genetically modifiable animal model such as *Drosophila* could allow for a better understanding of the role of Eiger/TNFα and other conserved cytokines in the FB or in the hemocytes. For example, using the combination of the double binary system (Yagi et al., 2010), a genetic tool commonly used to manipulate gene expression with spatial and temporal control in two different tissues, we could induce or suppress the expression of genes of interests in the FB or hemocytes at specific developmental stages to better understand the non-autonomous mechanisms underlying the migration of hemocytes responsible for chronic inflammation in obesity.

Activation of the IMD and Toll pathways in our OBL model is not a surprise, indeed, the IMD pathway is a key innate immune signaling pathway in *Drosophila,* like the mammalian NF-κB pathway. Activation of the IMD pathway occurs in response to microbial infection and stress, resulting in the production of antimicrobial peptides and immune effectors (Lemaitre and Hoffmann, 2007). In our OBL larvae, there is evidence to suggest that the activation of the IMD pathway is involved in triggering the immune response and hemocyte infiltration in the fat body. Similarly, the Toll pathway is another important immune signaling pathway in *Drosophila.* It plays a crucial role in defense against fungal and Gram-positive bacterial infections. Activation of the Toll pathway leads to the production of antimicrobial peptides and the induction of immune responses. In our context of obesity, the Toll pathway is also implicated in the immune activation and hemocyte infiltration in the fat body. The mechanisms by which the IMD and Toll pathways contribute to hemocyte infiltration in the fat body of obese larvae are not yet fully understood. However, it is believed that the immune activation and subsequent production of immune effectors and cytokines attract hemocytes to the fat body, leading to their infiltration and accumulation. Further research is needed to uncover the precise molecular mechanisms and downstream targets involved in these pathways during obesity-related immune responses in *Drosophila*.

### Final remarks

Several *Drosophila* models have been developed that exhibit features like obesity and metabolic disorders observed in humans. These models can be induced through genetic manipulations, high-calorie diets, or alterations in signaling pathways involved in metabolism and energy homeostasis. We believe we can use our OBL model to identify novel pathways or to screen for anti-obesity or anti diabetic drugs. As shown in Figure 4 (K-M), the administration of the anti-diabetes drug Metformin or the antioxidant anthocyanin significantly reduced the immune cell infiltration, demonstrating the potential utility of the model.

Our OBL model mimics the reduction of estrogen in females or testosterone in males, which has the effect of disrupting body fat distribution and is associated with greater risks of obesity-related metabolic disorders. In humans, obesity itself can also disrupt the hormonal balance in the body, including the regulation of steroid hormones (Fan et al., 2019; Sebo and Rodeheffer, 2021; Wittert and Grossmann, 2022). However, the relationship between steroid hormones, obesity, and metabolic regulation is complex and multifactorial. While steroid hormones play roles in both growth/maturation and metabolic regulation, the mechanisms underlying these processes are still being actively studied. Adipose tissue, both in flies and humans, also produces and secretes different factors including adipokines (TNFα and cytokines), which can influence various physiological processes, including metabolism and inflammation. These hormonal imbalances can further contribute to metabolic disturbances associated with obesity.

Studying these processes in model organisms like flies provides valuable insights into the fundamental mechanisms underlying growth and maturation in more complex organisms, including humans. By understanding these mechanisms, we can gain a better understanding of the regulation of growth and maturation in various species and potentially identify targets for therapeutic interventions related to growth and metabolic disorders.

We believe that our OBL model can be a valuable tool since it aims to mimic key features of human obesity and T2D, thus providing a platform either to perform genetic screens, or to test the efficacy and safety of novel therapeutic interventions. The models can be used to assess the impact of drugs on various metabolic parameters, including body weight, fat accumulation, glucose tolerance, insulin sensitivity, and lipid profiles. Additionally, the impact of drugs on adipose tissue metabolism and lipid accumulation can be examined. Drug screening in *Drosophila* allows for the rapid and cost-effective assessment of drug efficacy, toxicity, and potential side effects. The small size of the flies and the ability to perform high-throughput assays enable the screening of a large number of compounds. Moreover, the conservation of key metabolic pathways between Drosophila and mammals increases the likelihood of identifying compounds with potential relevance to human diseases.

## MATERIALS AND METHODS

### Fly stocks and husbandry

Fly cultures and crosses were raised at 25 °C, on a standard medium containing 9 g/L agar (ZN5 B and V), 75 g/L corn flour, 60 g/L white sugar, 30 g/L brewers’ yeast (Fisher Scientific), 50 g/L fresh yeast and 50 mL/L molasses (Naturitas), along with nipagin and propionic acid (Fisher). The lines were obtained *P0206-Gal4* and *UAS-NOC1-RNAi* (B25992)(Destefanis et al., 2022; Valenza et al., 2018), *HmlΔ-DsRed* (herein called *Hml-RFP*)(Makhijani et al., 2011), the lines mutant for *eiger^1^, eiger^3^, grnd, spzr^m7^, Rel^E20,^* and the reporter lines *Dpt-LacZ, Drs-GFP* were a gift of from Bruno Lemaitre (EPFL, Lausanne, CH); *eiger^1AG^* and *eiger^3AG^* and *eiger-GFP* (fTRG library, VDRC-318615) were a gift from Hugo Stocker (ETH, Zurich, CH).

### Hemocytes quantification

To label *in vivo* plasmatocytes, which comprise more than 95% of the hemocytes population in the *Drosophila* larva, we used the transgene line *Hml-Δ-Ds-RFP* (Makhijani et al., 2011). Fat bodies from larvae at 5 and 12 days AEL were dissected in phosphate-buffered saline (PBS) pH 7.4 and fixed in 4% paraformaldehyde (PFA) for 30 minutes at room temperature. Hoechst 33258 (Sigma Aldrich) was added to stain nuclei at the final concentration of 1 μg/ml. After washing with PBS, fat bodies were mounted onto slides with Vectashield (Vector Laboratories). Images were acquired using Leica-TCS-SP5 or TCS-SP8 confocal microscopes and the number of hemocytes and fat cells was counted from pictures taken at the same magnification.

### Immunofluorescence on hemocytes and IPC cells

Hemocytes in FB: dissected fat bodies from 20 larvae were fixed in 4% paraformaldehyde (PFA) (Electron Microscopy Science) in PBS for 40 minutes. After permeabilization with 0.3% Triton/PBS, tissues were washed in Tween 0.04% in PBS, saturated with 1% BSA in PBS, and incubated with anti-SPARC antibodies (1:400) (Martinek et al., 2002), overnight. Anti-Rabbit secondary Alexa555 was used 1:1000 (Invitrogen). After washing with PBST, fat bodies were mounted on slides using Vectashield (Vector Laboratories).

IPCs and anti-DILP2 immunostaining: larvae of the indicated genotype at 5 or 12 days AEL were either kept in normal fly food or starved for 24 hours in Petri dishes containing filter paper moistened with PBS. Brains were dissected in PBS and fixed in 4% PFA in PBS for 30 min and permeabilized in PBS-0.3% Triton X-100 (PBT) for 30 minutes. Tissues were blocked in 5% BSA in PBS and rat-anti-DILP2 was incubated overnight at 4°C (Geminard et al., 2009) followed by Alexa488 anti-rat-secondary antibodies (Invitrogen). Tissues were washed and mounted in Vectashield with DAPI (Vector Laboratories) and fluorescence images were acquired using a Leica-TCS-SP8 confocal microscope. The quantification of the mean pixels corresponding to the fluorescence intensity in the IPCs was determined using ImageJ, using similar parameters as previously described (Geminard et al., 2009; Parisi et al., 2013). Data are expressed in arbitrary units and represent the average fluorescence intensity in the IPCs of DILP2. Relative standard deviations were calculated using four to six independent experiments, each including at least ten animals of each genotype and condition.

### RNA extraction and quantitative RT-PCR analysis

Total RNA was extracted from 8 whole larvae, or from 10 dissected fat bodies using the QIAGEN RNeasy Mini Kit (Qiagen) according to the manufacturer’s instructions. Extracted RNAs were quantified using an ultraviolet (UV) spectrophotometer, and RNA integrity was confirmed with ethidium bromide staining. 1 μg total RNA from each genotype was reverse transcribed into cDNA using SuperScript IV MILO Master Mix (Invitrogen). The obtained cDNA was used as the template for quantitative real-time PCR (qRT-PCR) using qPCR Mastermix (Promega). mRNAs expression levels were normalized to *actin-5C mRNA* used as the internal control The relative level for each gene was calculated using the 2-DDCt method (Hulf et al., 2005) and reported as arbitrary units. Three independent experiments were performed and cDNAs used in triplicate.

The following primers were used for qRT-PCR:

*actin5c:* 5’ F-CAGATCATGTTCGAGACCTTCAAC;

5’ R-ACGACCGGAGGCGTACAG (Parisi et al., 2013).

*E74b*: 5’ F-GAATCCGTAGCCTCCGACTGT;

5’ R-AGGAGGGAGAGTGGTGGTGTT (Parisi et al., 2013).

*brummer*: 5’ F-ATATGGACCCCGTGTTTCAA; 5’ R-AGCTTGTCGTGCTCCGTTAT.

*FAS*: 5’ F-GTTGGGAGCGTGGTCTGTAT; 5’ R-GGTTTAGGCCAGCGTCAATA.

*Lsd2/Perilipin*: 5’ F-AGGAAGATAATGTGCCAGTTCCCG;

5’ R-GCTGCCACCAGACTGCTCCAC.

*eiger*: 5’ F-AAAGGTGGATGGCCTCACG; 5’-R TGCCGGTATGTGCATTGTTG.

*GstD1:* 5’ F-GACTCCCTGTACCCTAAGTGC; 5’ R-TCGGCTACGGTAAGGGAGTCA.

### Larval weight

Larvae of the indicated age and genotypes were weighed as a group of 20 for each genotype and age for 5 independent experiments

### Glucose, Glycogen and TAGs, and measurement

Ten whole larvae were sonicated for 5 seconds on ice in 200 µl of PBS for glucose assays, or in 0.1% PBS Tween for triglyceride assays, heat-inactivated at 70°C for 5 minutes to inactivate endogenous enzymes, before centrifugation at 4 C for 1 minute at 5000rpm. Supernatant was collected and centrifuged again at 4°C for 3 minutes at maximum speed and then used for the assays. Dissected fat bodies, ten-twenty, for each genotype and age, were placed in Eppendorf with 100ul ice-cold PBS, and sonicated following the same protocol as for whole larvae.

For Glucose assays, trehalose was converted to glucose by adding Trehalase (SigmaT8778) at a final concentration of 0.025 U/ml and incubating for 15 minutes at 37°C. The samples were mixed with 500 µl of Glucose Reagent HK (Sigma G3293) for 15 minutes at room temperature, and absorbance was measured at 340 nm according to the manufacturer’s protocol. Concentrations were determined by comparing absorbances to a standard curve generated with a glucose standard (Sigma G3285). Glucose measurements from undigested samples were subtracted to give the amount of glucose obtained from trehalose.

For hemolymph assays, hemolymph was pooled from 10 larvae and 1-2 ul collected, diluted 1:20 in PBS, and heat-inactivated at 70°C for 5 min. 2ul of diluted hemolymph was used for each glucose assay. Glycogen was measured by digesting each sample with 0.3U of Amyloglycosidase (Sigma A7420) for 3h at 37°C to convert glycogen to glucose. The samples were mixed with 500ul of Glucose Reagent HK (Sigma) for 15 minutes at room temperature, and absorbance was measured at 340nm according to the manufacturer’s protocol. Glucose measurements from undigested samples were subtracted to find the amount of glucose obtained from glycogen. A glucose standard from Sigma (G3285) was used to calculate sample concentrations.

Triglyceride measurements were performed by incubating 20 µl of sample with 400 µl of Free Glycerol Reagent (Sigma X) for 15 minutes at RT, followed by 100µl Triglyceride Reagent (Sigma X) for 15 minutes, and absorbance read at 540nm according to the manufacturer’s protocol. Concentrations were calculated by comparing absorbances to a standard curve generated using a triglyceride standard contained in the kit. Glucose and triglyceride concentrations were normalized against the total protein concentration in each sample as measured by a BCA assay (Pierce). Assays were repeated a minimum of three times.

### Lipidomic analysis

Fat bodies from Drosophila have been mechanically homogenized (IKA T25 Ultra-Turrax) in glass vials with chloroform/methanol 2:1 plus butylated hydroxytoluene (BHT) 0.01% as antioxidant. Lipids have been extracted after addition of KCl 0.05% plus a known amount of triheptadecanoin as internal standard. Vials have been shaken two hours at 4°C and the organic phase extracted after centrifugation at 2.500 rpm for 15 minutes. Later, a second extraction have been performed, the organic phases merged, dried in a stream of nitrogen, resuspended in chloroform/methanol 2:1 plus BHT and stored at −80°C in the dark until used. The aqueous phase has been dried, the resulting pellet dissolved with NaOH 0.1 N and the total protein amount quantified utilizing the bicinchoninic acid (BCA) assay. All the results have been normalized by the protein content.

### Quali- and quantitative analysis of fat bodies triglycerides

An aliquot of the lipid extract had been loaded onto a channeled Silica TLC plate (BioMap, Italy) which will be pre-run in hexane/diethyl ether/acetic acid 80:20:1 (v/v/v), heat-activated and developed up to 2.5 cm from the top. After run, the TLC plates have been sprayed with dichlorofluorescein (0.15% in ethanol) and, after drying in a stream of nitrogen, the silica spots corresponding to those of the known standards scraped off.

For TG determination, samples have been derivatized with methanol/HCl 3N for 20 minutes at 80°C and extracted with hexane/water prior gas-liquid chromatographic (GLC) analysis conducted on a DANI 1000 machine (DANI instruments, Milano, Italy) equipped with a flame ionization detector, an autosampler and a 30m, 0.32mm, 0.25 um MEGA-1 (Mega Columns, Legnano, Italy) fused silica column. GLC parameters have been as follows: flow of hydrogen at a constant pressure of 1 bar; injector temperature from 150 to 300°C; detector temperature 350°C; oven temperature ranging from 150 up to 330°C with appropriate gradients. The total run lasted 25 minutes. Chromatograms have been recorded and the area of each fatty acid peak quantified by the dedicated Clarity Software (Clarity, Prague, Czechia). The mass of TG has been evaluated after integration of the areas of the peaks of each fatty acid composing the lipid by subsequent sum and comparison with that of the internal standard. Data have been expressed as relative % (qualitative analysis of fatty acids) or by ug of single-total fatty acid/mg total protein.

### Cell size and Nile-Red staining

Fat bodies were dissected from larvae at the indicated days AEL, fixed for 30 min in 4% PFA in PBS, permeabilized 10 min in 0.2% Triton X-100 in PBS, and incubated for 30 min in a 0.01mM Nile Red Solution (SIGMA) and with 1:100 Alexa Fluor 488 Phalloidin to visualize the cytoskeleton through the binding between Phalloidin and F-actin, and with Hoechst 33258 for nuclei. After washing, the fat bodies were mounted in Vectashield with DAPI (Vector Laboratories), and fluorescence images were acquired using a Leica-TCS-SP5 confocal microscope, and the area of adipose cells for each fat body was calculated with ImageJ software.

### Measurement of the cardiomyocyte’s contraction

Larvae expressing the Hand-GFP reporter (Lo et al., 2007) that marks cardiomyocytes and pericardial cells, of the indicated age and genotypes, were embedded in a commercially available mounting putty to block their movement with the cardiac tube visible in their dorsal part (Fink et al., 2009). Movies of Hand-GFP contraction were taken and data elaborated using the Open-Source Physics (OPS) Software Tracker, Video Analysis and Modeling Tools (OPS) https://physlets.org/tracker/, at least 11-12 larvae for each genotype and age were used. Data were elaborated by measuring the number of contractions over time and p-value calculated using Student’s t-test analysis.

### Analysis of tGPH localization

Activation of insulin signaling was analyzed using the reporter line tGPH carrying the PH (pleckstrin homology) domain fused with green-fluorescent protein that translocates in the plasma membrane as a sensor of PI3K activation (Britton et al., 2002). Fat bodies were quickly dissected in artificial hemolymph (Fink et al., 2009) containing 108 mM NaCl2, 5 mM KCl, 2 mM CaCl2, 8 mM MgCl2, 1 mM NaH2PO4, 4 mM NaHCO3, 15 mM HEPES, 10 mM sucrose, and 5 mM trehalose, at pH 7.1, and placed in a 24 a well-plate at about 20 fats for each time point. 1mM final concentration of insulin (porcine from Sigma Aldrich) was added for the indicated time. After washing once with cold PBS, the fat bodies were fixed in 4% PFA and Hoechst 33258 (Sigma Aldrich) and mounted on the microscope slides using Vectashield (Vector laboratories). Images were acquired with a confocal microscope (Leica-TCS-SP5 and the localization of tGPH was analyzed in the cytoplasm or/and membrane from images of about 25-30 cells and quantified using ImageJ.

### Protein extractions and western blotting

The larval fat bodies (10 for each genotype) were dissected in Schneider’s medium serum-free and lysed in 100 μl of lysis buffer (50 mM Hepes/pH 7.4, 250 mM NaCl, 1 mM ethylenediaminetetraacetic acid (EDTA), 1.5% Triton X-100 containing a cocktail of phosphatases inhibitors (PhosSTOP 04906837001, Merck Life Science) and proteases inhibitors (Roche, cOmplete Merck Life Science). Samples were sonicated three times for 10 seconds using a Branson Ultrasonic Sonifier 250 (Branson, Darbury, CA, USA) equipped with a microtip set at 25% power. Tissue and cell debris were removed by centrifugation at 10,000× *g* for 30 min at 4 °C. Proteins in the crude extract were quantified by a bicinchoninic acid (BCA) Protein assay Reagent Kit (Pierce), following the manufacturer’s instructions with bovine serum albumin as the standard protein. For SDS-PAGE, samples were incubated for 8 min at 100 °C in standard reducing 1x loading buffer; 40 µg of total protein were run on a SDS-polyacrylamide gel and transferred onto nitrocellulose membranes (GE-Healthcare, Fisher Scientific Italia) After blocking in 5% (*w*/*v*) non-fat milk in tris-buffered saline (TBS)-0.05% Tween (TBS-T), membranes were incubated overnight with primary antibodies against *Drosophila* pAkt-Ser505 (1:500, #4054) from Cell Signaling, or Actin5c (1:200, #JL20) from Developmental Studies Hybridoma Bank (DSHB), University of Iowa, IA, USA. Appropriate secondary antibody, was incubated for 2 hours at room temperature, followed by washing. The signal was revealed with ChemiDoc Touch Imaging System (Bio-Rad Lab).

### In vivo detection of ROS using Dihydroethidium (DHE)

DHE was used to detect cytosolic superoxides and radical oxygen species (ROS). The reaction between DHE and superoxide anions generates a highly specific red fluorescent product (ethidium), which intercalates with DNA. ROS levels were detected in fat bodies as described in (Owusu-Ansah and Banerjee, 2009). Briefly, larval fat bodies at 5 and 12 days AEL were dissected in Schneider’s S2 medium (GIBCO). After incubation in 30 μM DHE (Invitrogen) for 5-7 minutes in the dark and at room temperature, fat bodies were washed three times with Schneider’s medium and immediately mounted with VECTASHIELD Antifade Mounting Medium. Images were taken using a Leica-TCS-SP5 confocal microscope.

### Treatment with Metformin, ACN, and HFD

Crosses were kept in culture bottles perforated to provide adequate air circulation for the parents and eggs were collected every 3 hours on a grape agar plate supplemented with life yeast. After 24 hours, first instar larvae were placed into vials containing standard corn meal (CM) alone or supplemented with the different chemical drugs: ACN: 0.24 mg/ml anthocyanins (gift from Katia Petroni and Chiara Tonelli, University of Milan) (Pilu et al., 2011). Metformin (Abcam) at 1mM final concentration, or in High Fat Diet (HFD) constituted of corn meal (CM) supplemented with 30% coconut oil (Trader Joe’s, Pasadena CA).

### Statistical analysis

The experiments were repeated at least three times and the statistical analysis among the various genotypes were examined by Student’s t-test or using One-way ANOVA with Tukey multi-comparisons, as indicated in the figure legends for each experiment, and calculated using GraphPad-PRISM 7 *p* values are indicated with asterisks **\* =** *p* < 0.05, ** =*p*< 0.01, *** =*p*< 0.001, **** =*p*< 0.0001, respectively.

## Acknowledgments

We thank you the Imaging Facility and CIBIO, and at the University of Milano. Miguele Ringuette University of Toronto (CA) for the anti-SPARC antibodies. Hugo Stocker at ETH (CH) for the anti DILP2 antibodies. Anthony Ferrante at Columbia University (USA) for the helpful discussion.

## Author contributions

Conceptualization: Z.M., A.V., S.Z., L.A., J.P., V.L., M.C., M.F., M.E.P., P.B.; Methodology, Validation and Formal Analysis: Z.M., A.V., S.Z., C.B., L.A., N.F., J.P., V.L., M.C., M.F., M.S., P.B.; Writing - original draft: Z.M., P.B.; Writing - review & editing: S.Z., P.B. L.A.; Supervision: P.B.; Project administration: P.B.; Funding acquisition: P.B.

