## supplementary for "A Novel Drosophila Model to Investigate Adipose Tissue Macrophage Infiltration (ATM) and Obesity highlights the Therapeutic Potential of Attenuating Eiger/TNFα Signaling to Ameliorate Insulin Resistance and ATM"

**Figure 1: larval growth and length and expression of *E74B-mRNA***

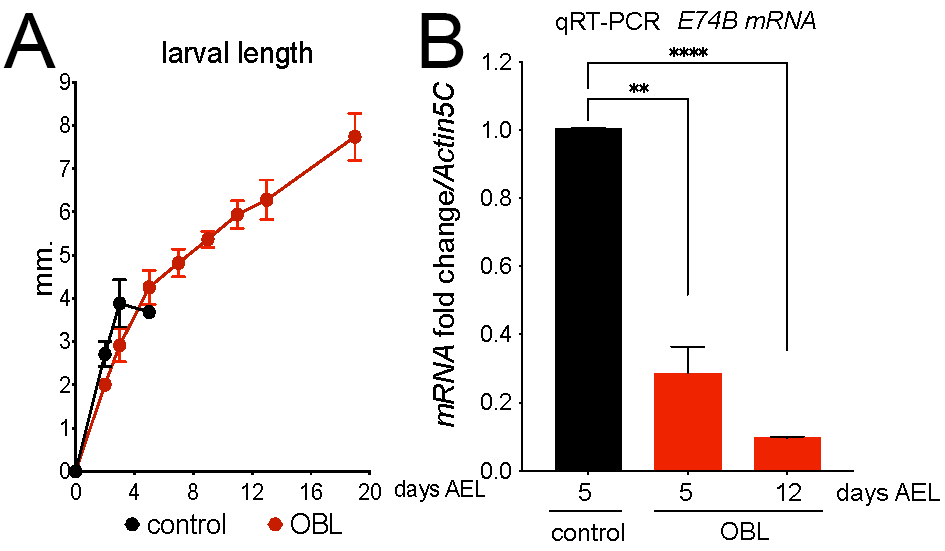

(A) Larval growth and length were measured over time of control *P0206>Gal4* or *P0206>Gal4; NOC1-RNAi* (OBL) larvae; *P0206>Gal4* larvae develop at the expected rate and between 5-6 days AEL undergo to metamorphosis, while OBL animals keep growing. (B) expression of *E74B.* To develop the OBL model we reduced the level of ecdysone in the prothoracic gland (PG), by reducing the expression of the *NOC1* gene using the NOC1-RNAi transgene under the control of the *P0206-Gal4* promoter (Valenza et al., 2018). Reduction of NOC1 results in a strong reduction of protein synthesis (Destefanis et al., 2022) which results in a decrease of Ecdysone production and circulation, shown by the decrease in the expression of its target E74B normally expressed in the FB. These animals, herein called OBL, *also show* a reduction of the size of their prothoracic gland (Destefanis et al., 2022), and because the level of ecdysone is low, they develop at a normal rate but never reach metamorphosis and continue to feed and gain weight until they reach an average of 20 days AEL before dying (Davis and Li, 2013; Quinn et al., 2012)

### Figure 2: Hemocytes analysis in various tissues of OBL larvae at 12 days AEL.

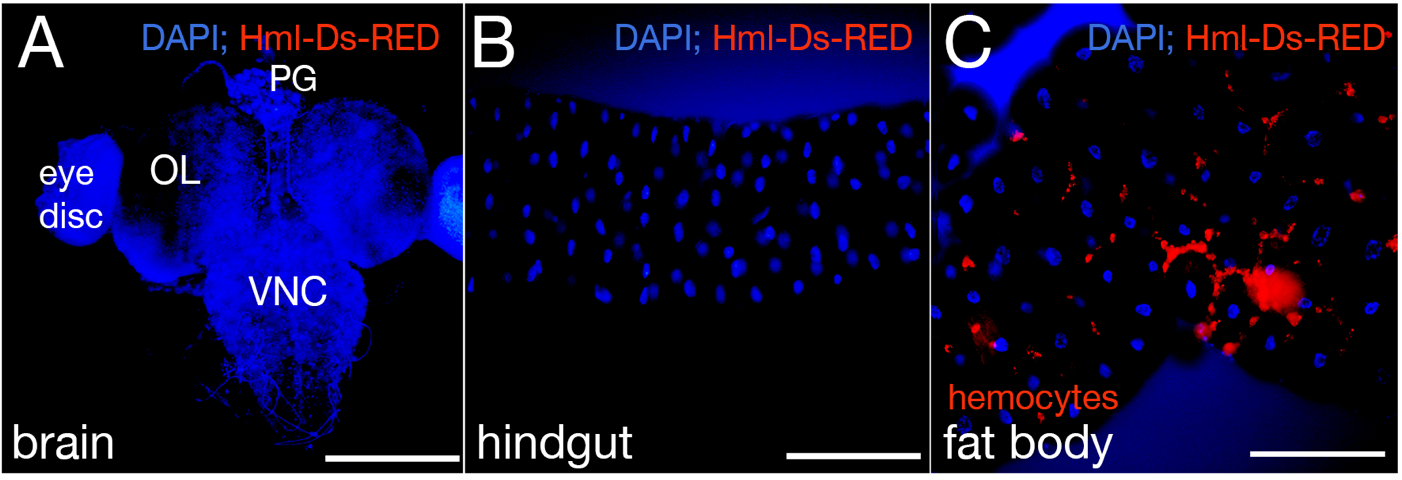

Confocal images of (A) brain, (B) hindgut, and (C) FB of larvae at 12 days AEL. Hemocytes are labeled in RED using the reporter line Hml-Ds-RFP (Makhijani et al., 2011), and the nuclei are stained with Hoechst in BLUE. VNC: Ventral Nerve Cord, OL: olfactory Lobe; PG, prothoracic gland. white bar represents 100 mm.

**Figure 3: Lipidomic analysis showing the Ratio of C14:0 myristate and C16:) palmitic Saturated Fatty Acids/Total lipids.**

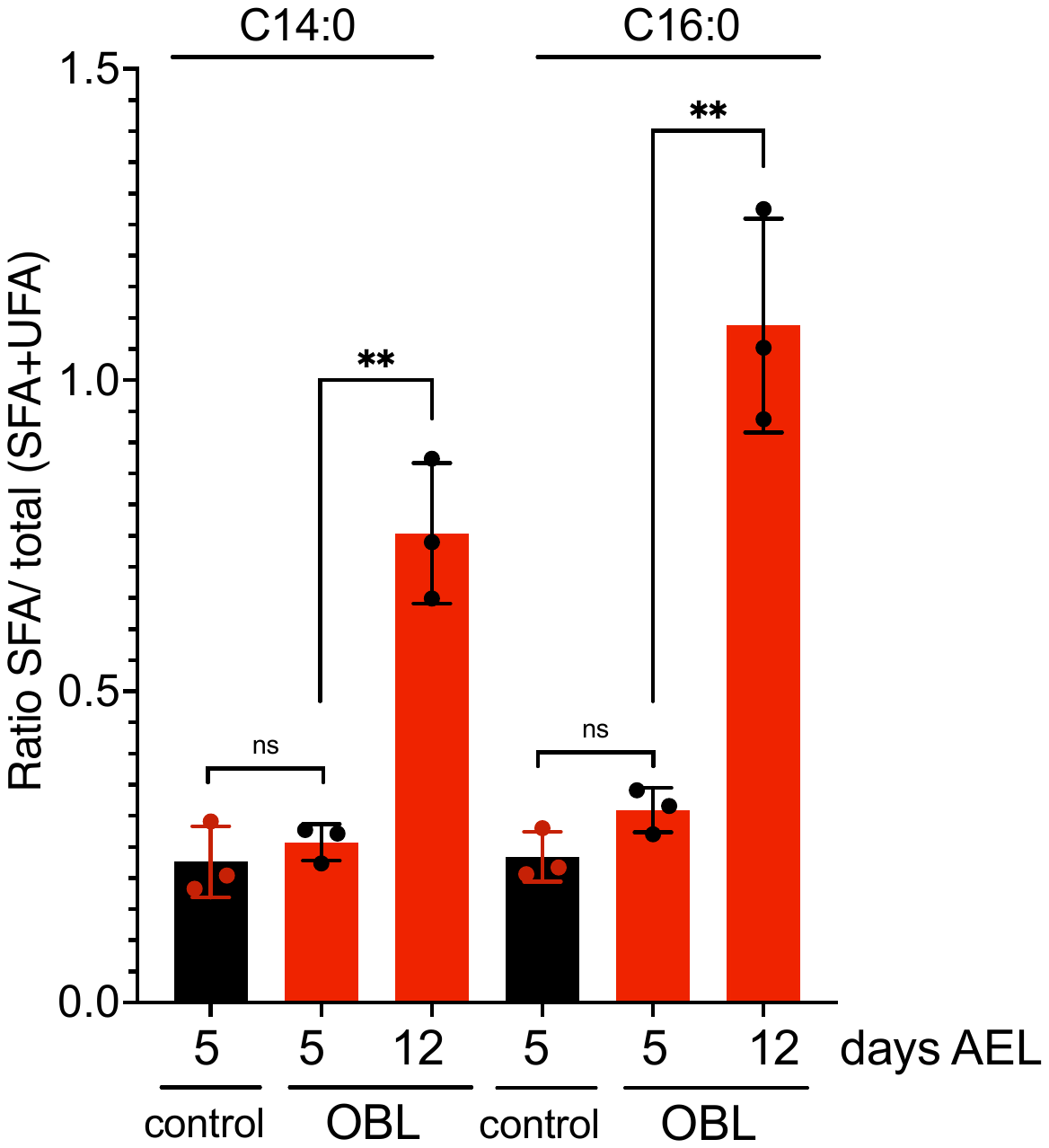

Lipidomic analysis was performed in control and OBL larvae at the indicated days AEL.

The graph represents the ratio of saturated fatty acids (SFA) over SFA plus unsaturated fatty acids (UFA). Statistical analysis was performed using one-way analysis of variance (ANOVA) with Tukey multiple comparisons. The asterisks represent the ** = *p* < 0.01, and the error bars indicate the standard deviations.

**Figure 4: Quantification of Glycogen and Glucose in whole larvae**

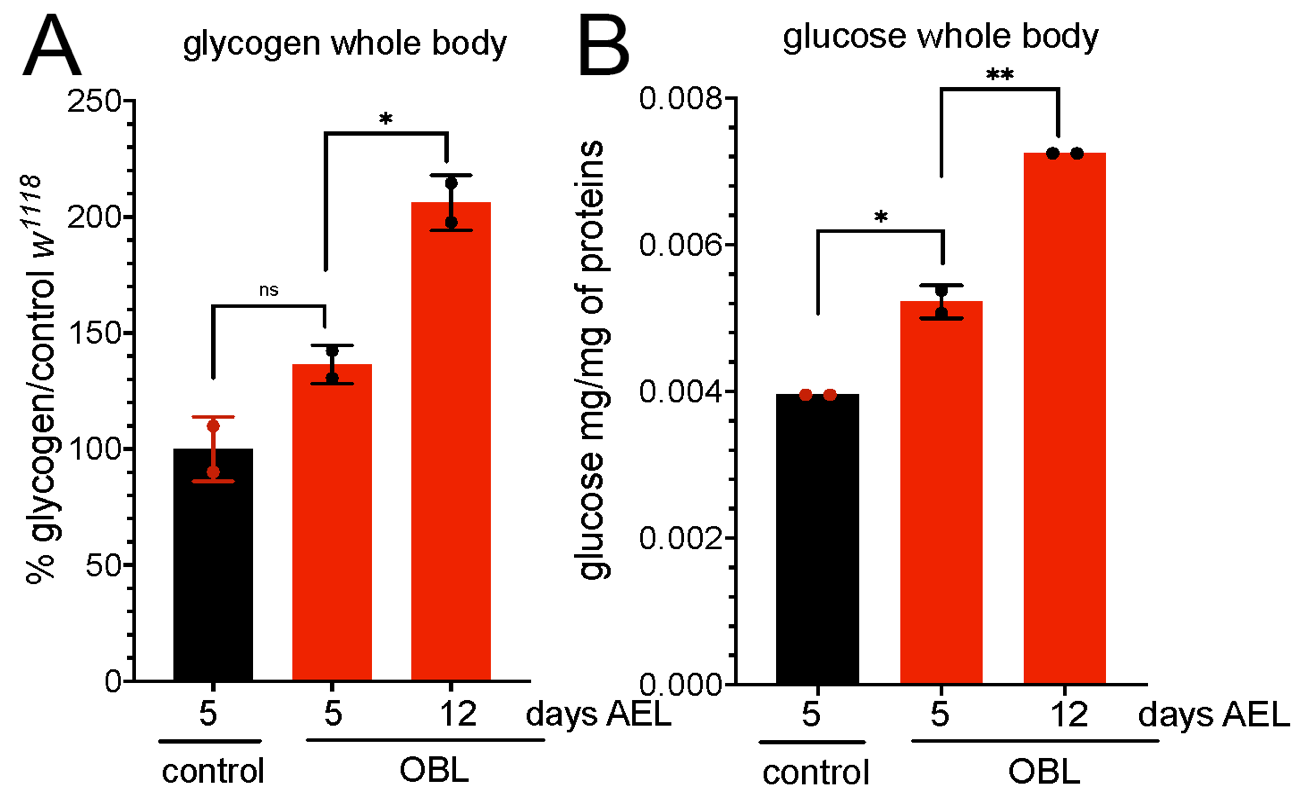

Quantification of glycogen (A) or glucose from threalose (B) was performed as described in material and methods using larvae of the indicated genotype and age. Statistical analysis was performed using Student’s t-test. The asterisks represent the * = *p* < 0.05, ** = *p* < 0.01, and the error bars indicate the standard deviations.

**Figure 5: Expression of *eiger-mRNA* and *GstD1-mRNA* in control and OBL animals**

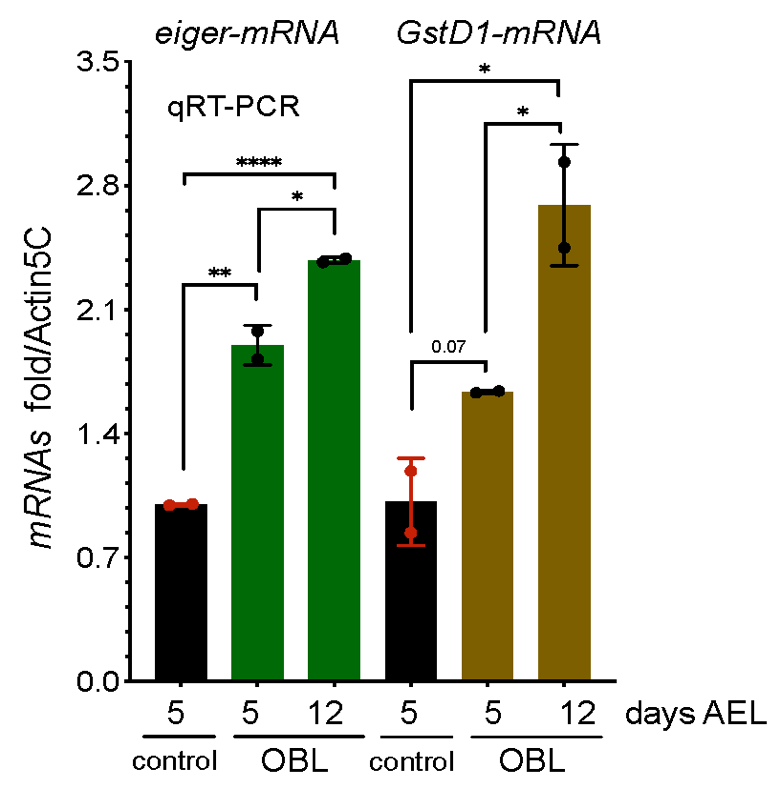

qRT-PCT showing the level of *eiger-mRNA* and of *GstD1-mRNAs* from whole larvae of the indicated genotype collected at 5 and 12 days AEL.
